# Recent and high grazing pressure limit cork oak seedling resprouting and survival

**DOI:** 10.1101/2025.11.20.689419

**Authors:** Abdullah Ibne Wadud, João Craveiro, Simone Erroi, Sandra Alcobia, Manuela Branco, Miguel N. Bugalho, Pedro Gonçalves Vaz

## Abstract

Regeneration failure is a bottleneck in Mediterranean oak woodlands. Cattle can hinder or promote recruitment, depending on grazing location, timing and intensity. Herbivory theory predicts that repeated defoliation and trampling deplete seedling reserves, whereas resprouting can extend survival; yet field studies rarely separate intensity from recency or combine long-run grazing records with individual fates and microhabitat/climate context. We test how management-driven heterogeneity shapes cork oak seedling survival and resprouting by combining 12 years of paddock-level grazing records with individual tracking of 8431 seedlings across 24 paddocks. Bayesian mixed-effects survival models related seedling lifespan to grazing history × pressure (moderate ≤150; high >150 LSU ha^-1^ days yr^-1^) and to key covariates, including seedling height, resprouting status, shrub distance, cattle dung counts (as a proxy of very recent grazing), and 1-month SPEI (as recent water balance). Bayesian logistic mixed models were then used to relate resprouting probability to grazing treatments. Survival was lower in grazed than ungrazed paddocks and declined along management gradients: median lifespan fell from 460 (moderate grazing) to 256 days (high), and from 460 (old grazing; two-year absence) to 199 days (recent). A two-year cattle absence increased survival under moderate pressure but was insufficient where pressure was high, indicating legacy effects and that recovery windows must scale with pressure. Resprouting dominated persistence: resprouters lived >5× longer than non-resprouters (2351 vs 460 days). Taller seedlings lived longer, and shrub proximity conferred a modest benefit. Climate modulated outcomes: wetter recent periods (higher SPEI) markedly boosted survival. Cattle reduced the odds of resprouting, with the strongest penalty under recent use. By disentangling grazing intensity from recency and linking both to seedling survival and resprouting, we show why recruitment falters under continuous, heavy grazing and when it can recover. Because drought intensifies cattle impacts, managers should combine moderate stocking rates with multi-year rest periods to rebuild oak bud banks and below-ground reserves; a two-year hiatus can help under moderate pressure but appears insufficient where pressure is high. Aligning rotational plans with drought outlooks and tracking simple field cues (seedling height, recent resprouting) offers a practical path to reconcile production with regeneration in Mediterranean wood-pastures.

**Highlights:**

- Twelve years of grazing records linked to 8431 cork oak seedling fates
- Recent grazing reduced survival and resprouting versus a two-year cattle absence
- High grazing shortened lifespan; two-year rest helped only under moderate pressure
- Resprouting was the strongest survival correlate; resprouters lived over 5× longer
- Wetter short-term water balance increased cork oak seedling longevity

## 1. Introduction

Regeneration is a critical bottleneck in oak woodlands (Pulido & Díaz, 2005; McEwan et al. 2011; Annighöfer et al., 2015). In Mediterranean oak agroforestry systems, seedling survival is shaped by intense biotic and abiotic filters such as herbivory, drought, and microsite limitation (Harper, 1977; Pulido & Díaz, 2005; Tyler et al., 2006; Vaz et al., 2019; Díaz et al., 2021). Herbivory theory predicts that repeated defoliation and trampling curtail photosynthetic area and deplete reserves, while resprouting can restore above-ground biomass after top-kill and thus extend survival (Bond & Midgley, 2001; Clarke et al., 2013; Milios et al., 2014; Mechergui et al., 2023). Yet, in working landscapes where livestock is central, grazing can either inhibit or facilitate recruitment depending on pressure, timing, and spatial distribution (López-Sánchez et al., 2014; Schieltz & Rubenstein, 2016). Clarifying when grazing depresses versus permits early establishment—and how resprouting mediates outcomes—remains a key ecological and management question.

Mediterranean oak woodlands, such as Portuguese cork oak (*Quercus suber*) montados and Spanish dehesas, are emblematic agrosilvopastoral systems that deliver substantial ecological and socio-economic benefits but face converging pressures from pathogens, herbivorous insects, wildfires, climate change, and management legacies (Branco & Ramos, 2009; Plieninger et al., 2015; Listopad et al., 2018; Vaz et al., 2013; Wadud et al., 2024). Cattle grazing is widespread globally and is a major land use in Iberian oak systems (Carmona et al., 2013). High grazing pressure can alter community structure, soil conditions, and regeneration dynamics (Bugalho et al., 2011; Filazzola et al., 2020; Vaz et al., 2024), whereas low to moderate use may open niches and reduce competitive pressure (Kuiters & Slim, 2003; Pulido & Díaz, 2005; McEvoy et al., 2006; Schieltz & Rubenstein, 2016). In rotational systems, this interplay creates strong spatiotemporal heterogeneity in both grazing pressure and grazing history—features likely to influence seedling resprouting and survival but seldom quantified jointly with multi-year management records and individual seedling fates (López-Sánchez et al., 2014).

Resprouting is a key persistence mechanism for woody plants exposed to recurrent disturbance, allowing seedlings to replace lost shoots after browsing or top-kill and thus extend survival under grazing pressure. In Mediterranean oaks, including *Q. suber*, this capacity depends on protected basal buds and below-ground reserves, but repeated defoliation can progressively deplete those reserves and limit recovery under sustained grazing (Arosa et al., 2015; Catry et al., 2010; Monfort-Bague et al., 2020; Mechergui et al., 2023). This suggests that resprouting may buffer early damage, yet its effectiveness should decline where grazing is more intense or more recent.

Several proximate covariates govern early performance. Seedling size (height) integrates reserves and rooting depth, predicting tolerance to damage and drought (Tyler et al., 2006; Clarke et al., 2013; Vaz et al., 2019). Under continued grazing, greater seedling height may also allow partial escape from browsing and improve persistence (Uytvanck et al., 2010). Shrubs can act as nurse cover or, conversely, intensify competition, with consequences for microsite moisture and protection from trampling (Pulido & Díaz, 2005; Díaz et al., 2021; Listopad et al., 2018). Finally, short-term climatic water balance modulates hydraulic stress during vulnerable stages, potentially altering survival under otherwise similar grazing regimes (Díaz et al., 2021). Against this background, robust inference demands designs that couple long-run grazing information with individual-level fates and microhabitat/climate context in real paddocks.

Using a previously unpublished 12-year, paddock-level dataset on cattle grazing pressure under rotational grazing, together with individual survival monitoring of 8431 cork oak seedlings, we tested how cattle grazing alters cork oak regeneration—seedling survival and resprouting. First, we tested how seedling survival differs between ungrazed and grazed paddocks, predicting higher survival in the absence of cattle. Second, within grazed paddocks, we evaluated how grazing pressure (moderate vs. high) and grazing history (old, i.e. two-year cattle absence, vs. recent) affect survival, expecting lower survival under high pressure and under recent grazing. Because successful resprouting after top-kill depends on protected buds and below-ground reserves, we expected seedlings observed to resprout during the study to persist longer than those not observed to resprout. Third, we examined whether cattle grazing reduces the probability that seedlings resprout and whether resprouting probability varies with grazing pressure and history, predicting lower resprouting under high pressure and recent grazing.

## 2. Materials and methods

### 2.1. Study area

The study was conducted between November 2021 and February 2023 in an 18000-ha cork oak woodland—known as *montado* (or *dehesa* in Spain)—in central Portugal (Fig. 1). The area comprises fenced paddocks, some grazed and others ungrazed, enclosed by barbed-wire fencing, and is part of Companhia das Lezírias (CL; 38°50′ N, 8°49′ W), a multi-use, state-owned agrosilvopastoral estate and core site of the Long-Term Socio-Ecological Research platform Montado (www.ltsermontado.pt). Since the 19th century, CL has been managed for cork production, extensive cattle grazing, recreational hunting, and, more recently, biodiversity conservation. The region has a Mediterranean climate with hot, dry summers and mild, wet winters (Craveiro et al., 2019; Azedo et al., 2022). In 2022, annual precipitation was 502 mm, and the mean monthly temperature was 17.1 °C (min/max: 5.6–30.8 °C). The terrain is mostly flat (1–53 m elevation), with sandy, acidic soils (pH 5.4–6.3) low in organic matter and water retention, predominantly Podzols and hydromorphic types (Concostrina-Zubiri et al., 2017).

**Figure 1.**
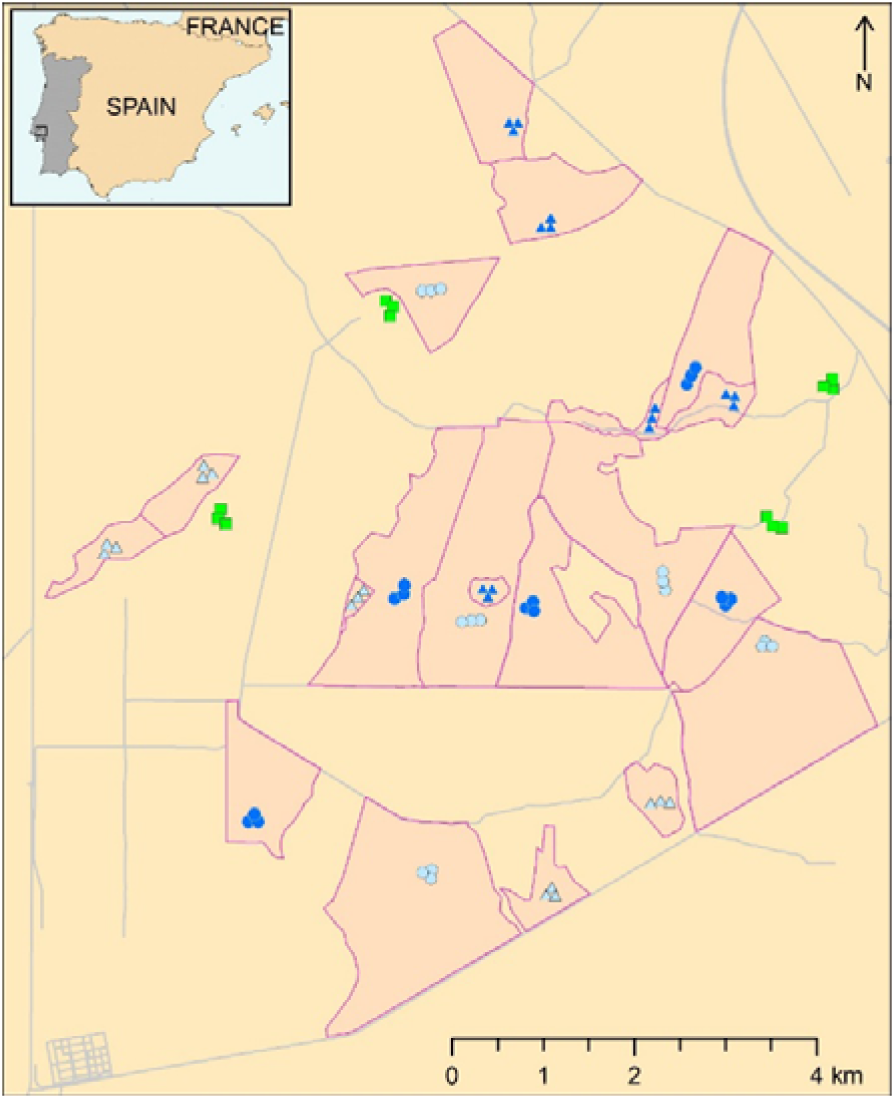
Study area at Companhia das Lezírias (central Portugal) and sampling design. Paddock boundaries are shown in pink. Symbols mark the three sampled adult cork-oak trees per paddock (20-m radius plots) where all naturally established *Quercus suber* seedlings (≤15 cm) were tagged and monitored. Color denotes grazing pressure (dark blue = high; light blue = moderate; 12-year records) and shape denotes grazing history (triangles = old, two-year cattle absence prior to the study; circles = recent, grazed in the previous two years). Green squares indicate the sampled trees in long-term ungrazed paddocks (>12 years; paddock polygons not shown for ungrazed).

Cork oak trees at CL have a mean basal area of 13.0 m²/ha and an average density of 53.6 trees/ha (Mexia et al., 2022), with 24–49% canopy cover (Listopad et al., 2018). Most individuals are 4–14 m tall and 10–45 cm in diameter at breast height, DBH (Vaz et al., 2011). The most common understory species are *Ulex australis*, *Cistus salviifolius*, *Myrtus communis*, *Cistus crispus*, *Genista triacanthus*, and *Cistus ladanifer* (Carrilho et al., 2017), along with annual grasses and forbs such as *Tolpis barbata*, *Agrostis pourretii*, and *Ornithopus compressus* (Vaz et al., 2024). Cattle grazing is extensive and rotational, with herds of 50–300 head moved among paddocks. Stocking rates range from 0.5 to 1.1 LSU ha^-1^, typically lower in summer (LSU = livestock unit, a standardized adult-cattle equivalent used to express grazing load) (Gonçalves et al., 2012).

### 2.2. Study design

We used 12 years of grazing records (2009–2021) to classify paddocks by grazing pressure and history. None of the selected paddocks burned during this 12-year period or during seedling monitoring. All paddocks were pre-existing fenced units within the study area; in grazed paddocks, cattle were rotationally moved among them. All paddocks belonged to the same estate and, where grazed, were managed under the same overall extensive rotational grazing system. Grazing pressure was classified as moderate (≤150 LSU ha^-1^ days year^-1^) or high (>150), following thresholds used in regional grazing studies and chosen to maximize balanced sampling across paddocks and maintain a management-relevant factorial design (Gonçalves et al., 2012; Arosa et al., 2017). Paddocks with consistently low grazing were not included due to limited replication. Grazing history in grazed paddocks was categorized as recent (cattle present during the two years prior to and typically during the study) or old (no cattle for two years before the study), although all had been grazed between 2009 and 2019.

To test the effects of grazing pressure and history on seedling survival, we selected 20 grazed paddocks—five per treatment combination: old–moderate, old–high, recent–moderate, and recent–high. For the grazed vs. ungrazed comparison, we selected four of these grazed paddocks (one per treatment) and matched them with four nearby ungrazed paddocks (>12 years without cattle) with similar habitat. Paddocks varied in area (Fig. 1), but grazing pressure was quantified as LSU ha^-1^ days yr^-1^, standardizing for area and time.

### 2.3. Data collection

In each paddock, we selected three healthy adult cork oaks (10–45 cm DBH) using a haphazard but constrained procedure. We selected the first tree >50 m from paddock edges, and then two additional trees further within the paddock, each at least 100 m from the others. All naturally established cork oak seedlings ≤15 cm tall within a 20-m-radius plot around each selected tree were tagged with numbered plastic binder spines, and their positions recorded using polar coordinates—distance to the target tree (measured with a laser rangefinder, 1 mm precision) and azimuth angle (measured with a sighting compass from magnetic north). We did not determine the exact establishment date of individual seedlings; therefore, the tagged cohort may have included seedlings established at different times before the first census. In *Q. suber*, seedling establishment occurs predominantly through acorn germination, whereas vegetative resprouting is more typically basal in larger individuals, arising from the root collar or trunk base after severe damage (Catry et al., 2017, 2022), although we could not completely exclude occasional vegetative origin.

Seedling survival was monitored seven times (Nov–Dec 2021; Jan–Feb, Mar, Jun–Jul, Aug, Sep 2022; Feb 2023), with census intervals adapted to field logistics and seasonal conditions. Only seedlings recorded at the first census were included; new recruits were negligible during the sampling period and were excluded from analysis. Resprouting was noted as a binary variable (yes/no). A resprouting seedling was defined as a tagged seedling that showed reemergence of green tissue after having been previously fully desiccated or with no visible aboveground biomass during the monitoring period. A non-resprouting seedling was a tagged seedling for which no such reemergence was observed during monitoring. Because only seedlings with visible aboveground tissue at the first census could be tagged, individuals that had already lost all aboveground biomass before that census and resprouted later were not included. Seedlings were considered dead if they failed to resprout within 384 days (Vaz et al., 2019), remained unchanged across consecutive censuses, or were found uprooted with tags attached or nearby. Because all tagged seedlings were monitored for more than 384 days from the first census to the final census, this criterion could be applied within the study period.

Seedling height (main stem, in mm) was measured during the first, fourth, and final census. For seedlings that died earlier, the last recorded value was used. Microhabitat was characterized by the distance to the edge of the tree crown (not used in analyses, as nearly all seedlings were under canopy) and distance to the nearest shrub canopy edge (positive values indicating open ground).

To capture localized grazing intensity, cattle dung was counted within each 20-m-radius plot centered on the focal tree at all seven censuses, while avoiding re-counting pats already recorded in previous censuses. For each seedling, we used the dung count from its corresponding tree at each census or the last recorded value if it died earlier. In old grazing paddocks (two-year cattle absence), dung counts were typically near zero and therefore capture any occasional incursions; in recent grazing paddocks they track spatial variation in current use.

To characterize recent climate, we calculated the Standardized Precipitation–Evapotranspiration Index (SPEI) at a 1-month scale using the *SPEI* v1.7 R package (Vicente-Serrano et al., 2010; Beguería & Vicente-Serrano, 2023). Monthly climate data from CL (Jan 2021–Feb 2023) included precipitation, minimum and maximum air temperature, and extraterrestrial radiation (Ra). Thus, the SPEI calculation included monthly climate data from the period preceding the first census in Nov–Dec 2021. Potential evapotranspiration (PET) was estimated using the Hargreaves method (Tmin, Tmax, Ra, latitude). Climatic water balance was computed as precipitation minus PET, from which SPEI was derived; SPEI values preceding each census were then included in the survival models. Positive SPEI values indicate wet conditions, and negative values indicate drought, relative to the climatic series used for standardization.

### 2.4. Statistical analysis

#### 2.4.1. Seedling survival

To test whether seedling survival differed between grazed and ungrazed paddocks, we fitted Bayesian mixed-effects survival models assuming alternative response distributions (lognormal, Weibull, and exponential), with the lognormal distribution ultimately retained based on leave-one-out cross-validation. Survival time (days since first census) was modeled as interval-censored when death occurred between censuses and right-censored if the seedling was still alive at study end. Grazing treatment (grazed vs. ungrazed) was included as a fixed effect, and random intercepts accounted for seedlings nested within trees and trees within paddocks.

To assess effects of grazing pressure and history, we fitted a second survival model with the same censoring structure and compared the same alternative response distributions, again retaining the lognormal form. Fixed effects included grazing pressure (moderate vs. high), grazing history (old vs. recent), and their interaction. We added five covariates: seedling height, resprouting status (yes/no), distance to the nearest shrub, number of dung piles (proxy for localized grazing), and SPEI. All continuous predictors were centered and standardized; variance inflation factors were <2.1, indicating low multicollinearity (Lüdecke et al., 2021). For parsimony, we also fitted an additive model (without interaction) but leave-one-out cross-validation of expected log predictive density (ΔELPD LOO) favored the interaction model (ΔELPD LOO = −3.6 ± 2.5 SE; Vehtari et al., 2017). We further tested the removal of each covariate (excluding grazing variables) one at a time, interpreting ΔELPD LOO differences >2 × SE as meaningful (Vaz et al., 2021); results supported the full model.

Models were fitted using the *brms* package (Bürkner, 2017) in Stan (http://mc-stan.org/). Weakly informative priors were used to improve convergence while avoiding overfitting (Gelman et al., 2020): Normal(0, 3) for the fixed effect in the grazed vs. ungrazed model, and Normal(0, 5) and Normal(0, 1) for categorical and numeric predictors, respectively, in the grazing pressure × history model. Each model was run with four MCMC chains (4000 iterations, 1000 warm-up), yielding 12000 post-warm-up samples. Convergence was assessed via trace plots and RC (all ≤1.1). Leave-one-out cross-validation indicated that the lognormal distribution consistently outperformed the Weibull and exponential alternatives (lower LOOIC). Posterior predictive checks confirmed model adequacy.

#### 2.4.2. Resprouting probability

To assess whether grazing influenced seedling resprouting (yes/no), we fitted two Bayesian mixed-effects logistic regression models (Bernoulli, logit link). The first included grazing treatment (grazed vs. ungrazed) as a fixed effect, with a Normal(0, 4) prior. The second included grazing pressure and history, with Normal(0, 8) priors. Both models accounted for trees nested within paddocks in the random-effects structure. For the second model, we tested the interaction between grazing pressure and history but found no support for improved fit (ΔELPD LOO = −0.2 ± 0.5 SE), and thus retained the additive model. Diagnostics, convergence checks, and inference procedures followed those described above.

All analyses were conducted in R v.4.5.0 (R Core Team, 2024). Inference was based on posterior distributions and model predictions, with 95% credible intervals reported as summaries of posterior uncertainty (McElreath, 2018).

## 3. Results

We monitored a total of 8431 cork oak seedlings across the 24 paddocks. By the end of the study period, 53% of the seedlings had survived (Table 1, Fig. 2). Most seedlings (99%) could be identified under the canopy of adult cork oaks at the final census, while 15% were located beneath shrub cover. Survival was higher in the four ungrazed paddocks, where 62% of 1051 individuals persisted, compared to 51% survival among the 7380 seedlings in grazed paddocks. Within grazed areas, 58% of 4628 seedlings survived in paddocks with older grazing history—where cattle had been absent during the last two years—compared to 39% of 2752 seedlings in paddocks with recent grazing activity. Under moderate grazing pressure, 52% of 5679 seedlings survived, compared to 49% of 1701 under high pressure. Among the 4441 seedlings that survived, 30% were observed to resprout at least once during monitoring. By contrast, among the 3990 seedlings that ultimately died, only 1% were observed to resprout before dying.

**Figure 2.**
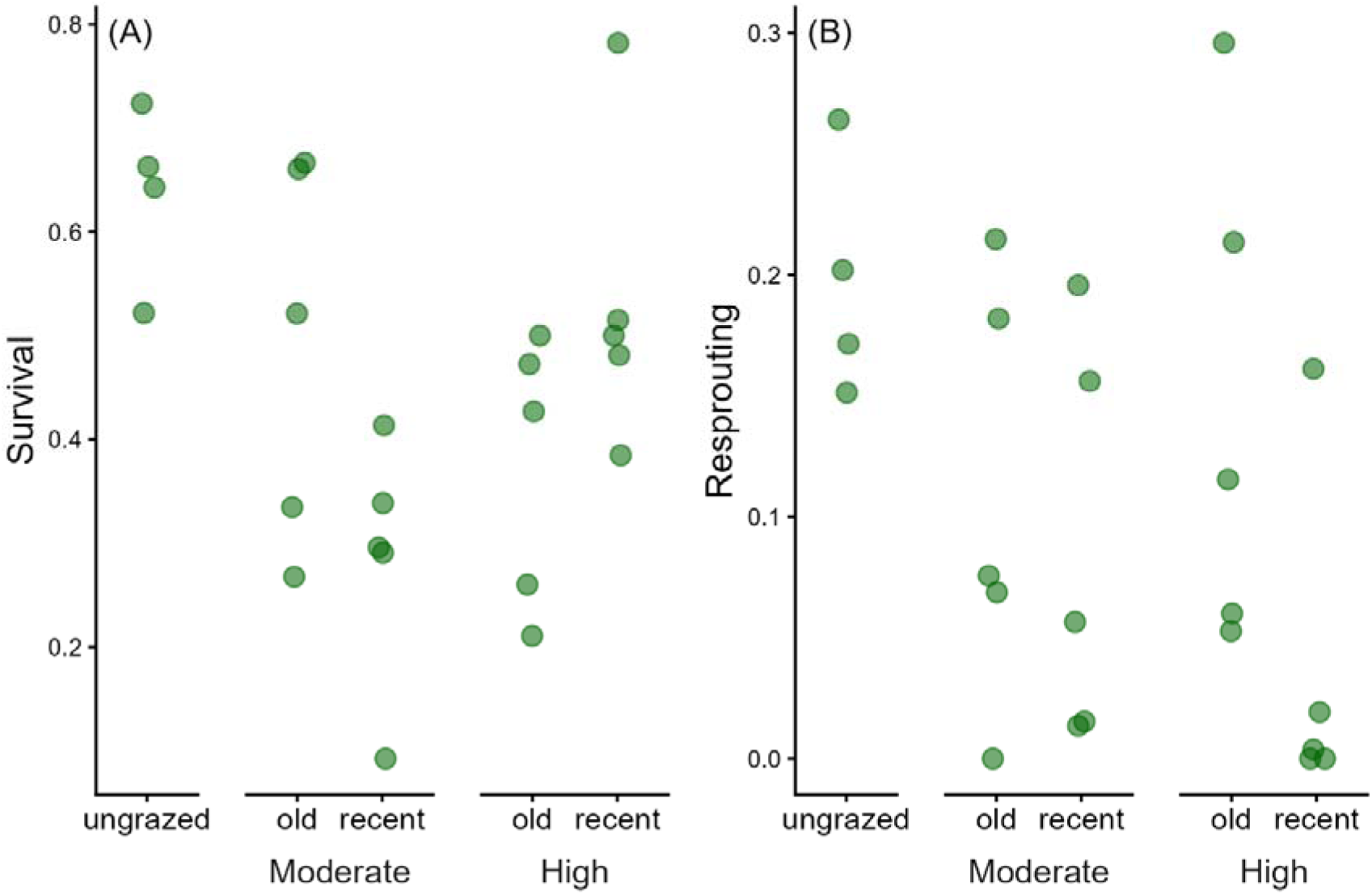
Raw seedling survival and resprouting across grazing treatments. Circles show each paddock’s observed proportion of seedlings that (A) survived to the final census and (B) resprouted at least once during monitoring. Grazed paddocks are grouped by grazing pressure (moderate vs high) and grazing history (old = two-year cattle absence prior to the study; recent = cattle present in the two years prior to and typically during the study). Ungrazed paddocks had been without cattle for >12 years.

**Table 1.**
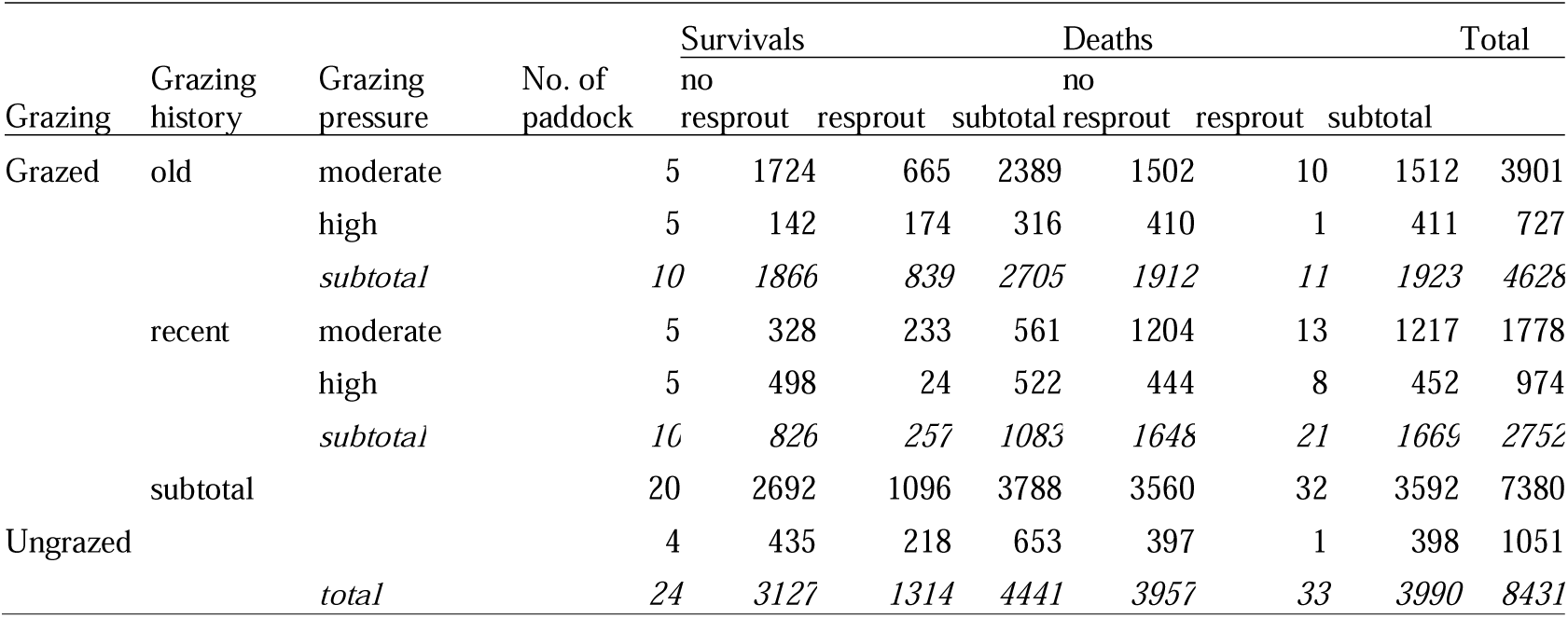
Number of cork oak seedlings that survived, resprouted, and died in each grazing treatment. Seedling survival was monitored within 24 paddocks distributed as shown. Within each paddock, survival was monitored around (20-m circles) 3 cork oak trees.

### 3.1. Survival: grazed vs. ungrazed paddocks

After accounting for random effects (trees nested within paddocks), our Bayesian mixed-effects lognormal survival model (Table A.1, Appendix A) indicated that seedlings in the four ungrazed paddocks had higher survival than those in four nearby grazed paddocks with similar conditions. We estimate a 98% posterior probability that seedling survival was higher in the absence of cattle grazing, showing strong support for this contrast (Bürkner, 2017; Clark, 2020; Vaz et al., 2021). The model-predicted median survival time was 302 days (95% credible interval: 226–421) in grazed paddocks, compared to 493 days in ungrazed paddocks—a 63% increase (95% CI: 363–653) (Fig. 3).

**Figure 3.**
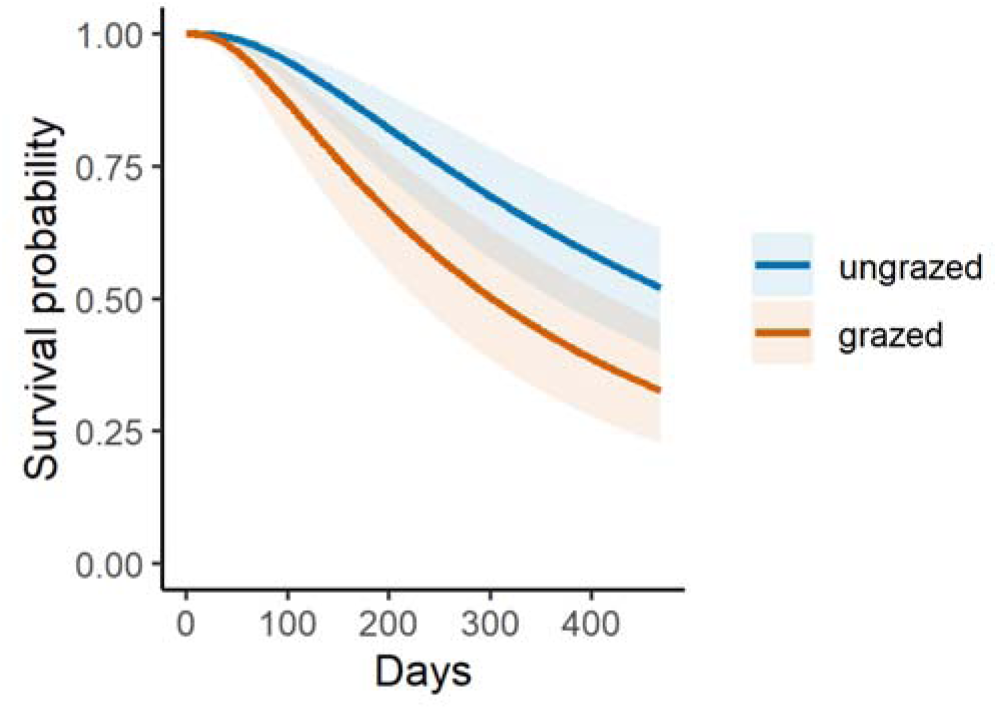
Fitted survival curves (± 95% credible intervals) showing the probability that tagged cork oak seedlings remained alive through time in ungrazed and cattle-grazed paddocks, estimated using a Bayesian mixed-effects lognormal survival model.

### 3.2. Survival: grazing pressure and grazing history

The Bayesian survival model (Table A.1, Appendix A) revealed that both grazing pressure and grazing history affected seedling longevity. Median predicted survival time was lower under high grazing pressure (256 days) than under moderate pressure (460 days; Fig. 4A). Similarly, seedlings in paddocks with recent grazing history had shorter predicted survival (199 days) than those with older grazing history (460 days; Fig. 4B). We found strong posterior support for these effects: there was a 97% probability that high grazing pressure reduced survival compared to moderate pressure, and a 100% probability that recent grazing had a negative effect compared to older grazing.

**Figure 4.**
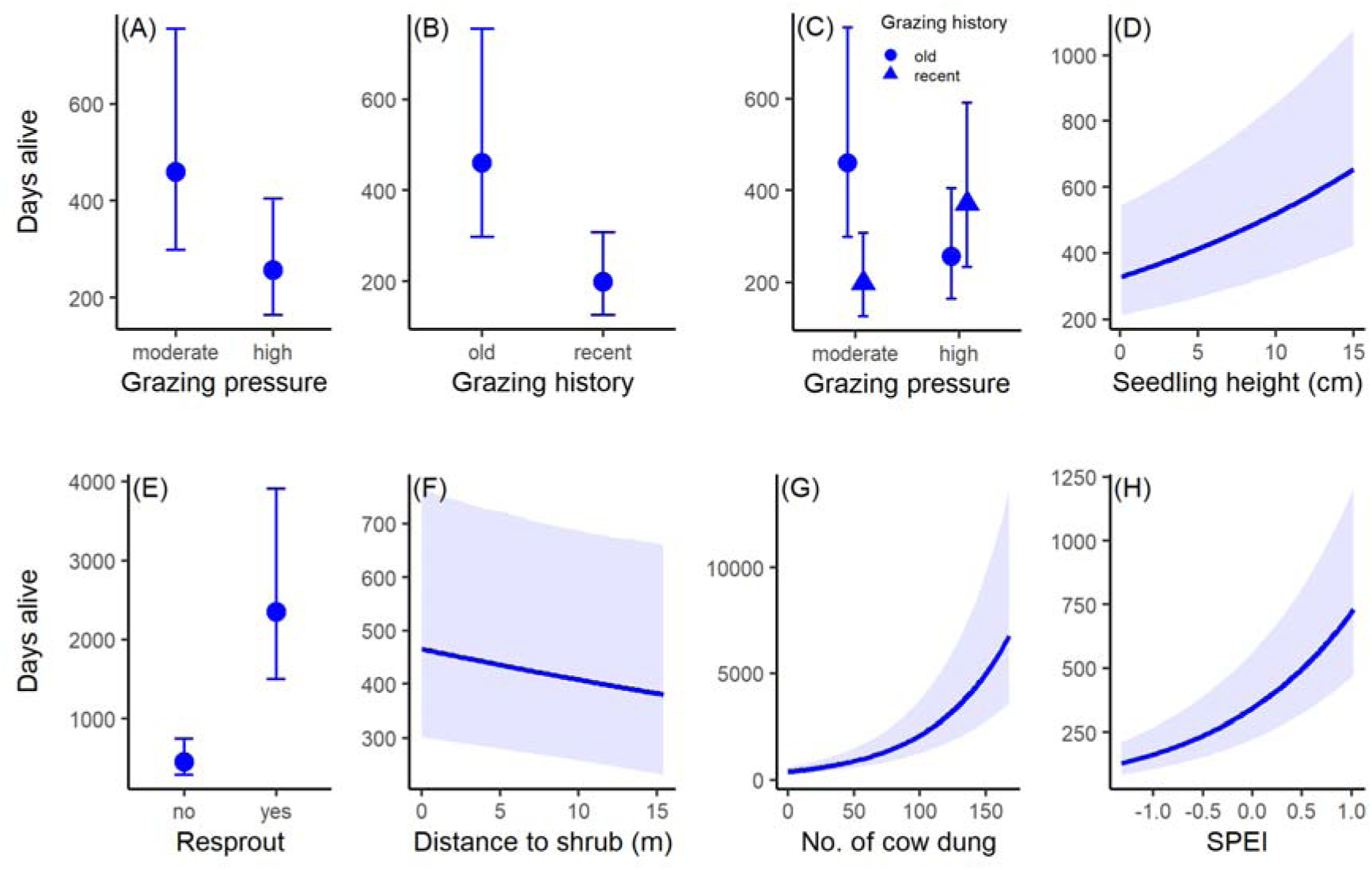
Model-fitted survival times for cork oak seedlings as a function of grazing treatments and covariates, based on the Bayesian mixed-effects lognormal survival model. Panels show effects of grazing pressure, grazing history, their interaction, and additional covariates. Points or lines represent posterior medians; shaded areas and vertical bars indicate 95% credible intervals. See main text for panel descriptions.

Analysis of the interaction between grazing pressure and grazing history (Fig. 4C) showed that the effect of grazing history differed between grazing pressure levels. Under moderate pressure, posterior support favored greater survival in the old than in the recent grazing condition (80% of posterior draws). Under high pressure, the predicted contrast was weak and reversed in direction, with recent grazing associated with slightly higher expected survival than old grazing, but uncertainty around this contrast was large.

Among the covariates, seedling height and resprouting status were positively associated with survival. The model predicts that a 15 cm seedling has a 58% longer lifespan than a 5 cm seedling (654 vs. 413 days). Successful resprouting was associated with the greatest increase in predicted persistence among all predictors: seedlings observed to resprout during the study persisted much longer than seedlings not observed to resprout, reaching a predicted lifespan of 2351 versus 460 days (Fig. 4E). In contrast, distance from shrub cover had the weakest effect. Still, there was a 95% probability that seedlings located farther from shrubs had slightly lower survival. For example, seedlings located 5 m from a shrub had a 6% shorter predicted lifespan than those beneath shrub cover (436 vs. 466 days; Fig. 4F).

Two additional covariates, cattle dung abundance and SPEI, were positively associated with seedling survival. Increased dung counts, used as a proxy for recent and localized cattle presence, were linked to longer survival times, with model predictions indicating a twofold increase in survival between areas with 50 and 100 dung piles (Fig. 4G). This pattern may reflect favourable microsites and reduced browsing near fresh excreta. Higher SPEI values, indicative of wetter recent conditions, were also associated with increased survival (Fig. 4H).

### 3.3. Resprout: grazed vs. ungrazed paddocks

Resprouting was more likely among tagged seedlings in ungrazed paddocks than among tagged seedlings in grazed paddocks. According to our Bayesian mixed-effects logistic regression model (Table A.2, Appendix A), the posterior probability of resprouting was 0.19 (95% CI: 0.13–0.26) in ungrazed paddocks, compared to 0.10 (0.05–0.17) in grazed paddocks (a 47% decrease; Fig. 5A). The model estimated a negative effect of grazing on resprouting, with 98% of posterior draws supporting a lower probability in grazed paddocks.

**Figure 5.**
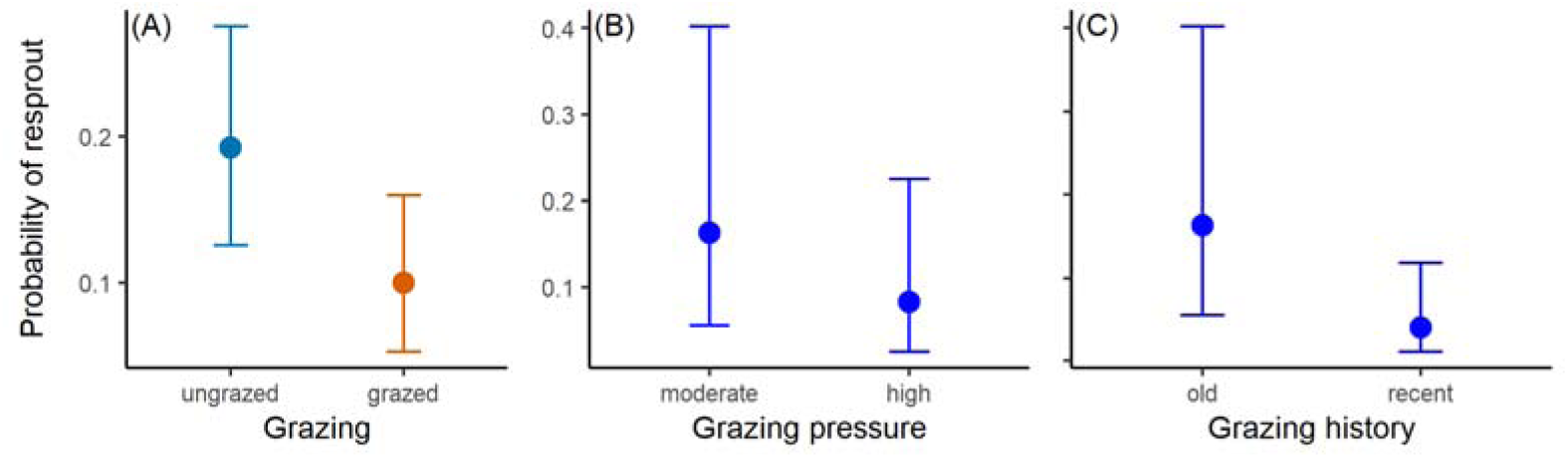
Model-fitted probability of resprouting in cork oak seedlings (posterior median ± 95% credible intervals) in response to grazing treatments, based on Bayesian mixed-effects logistic regression models. See main text for panel descriptions.

### 3.4. Resprout: grazing pressure and grazing history

Among tagged seedlings, resprouting was half as likely under high grazing pressure as under moderate pressure (0.08 vs. 0.16; Fig. 5B). Overall, 87% of posterior draws supported reduced resprouting under high grazing. In contrast, there was strong evidence for a negative effect of recent grazing history, with 99% of posterior draws supporting lower resprouting probability. Seedlings in recently grazed paddocks had a resprouting probability of 0.04, one quarter of that observed in paddocks with older grazing history (0.16; Fig. 5C).

## 4. Discussion

Our data provide management-relevant field evidence on a long-standing question in Mediterranean woodlands: how cattle use shapes the fate of cork oak seedlings and, more broadly, of young trees establishing under recurrent disturbance. Leveraging 12 years of paddock-level grazing records and individual monitoring of 8431 seedlings, together with microhabitat and short-term climatic water balance (SPEI), we disentangle grazing intensity from recency and link them to both survival and resprouting capacity. We discuss these results and their implications for the management of Mediterranean wood-pastures.

### 4.1. Grazing and resprout effects on seedling survival

Seedling survival was higher in ungrazed than in grazed paddocks, echoing patterns for Mediterranean oaks in Iberian agroforestry systems (Plieninger et al., 2010; Ritsche et al., 2021). The net negative signal of grazing accords with classic herbivory mechanisms—direct shoot consumption, repeated nipping and trampling, and soil compaction—plus shifts in soil structure and water infiltration that constrain root establishment (Pulido et al., 2010; Laskurain et al., 2013; Moradi et al., 2021; Palmerlee et al., 2025). Although grazing can suppress annual grasses and overall cover, potentially easing competition and sometimes reducing pressure from insects or small mammals (Tyler et al., 2008; Muñoz et al., 2009; Canelo et al., 2025; Vaz et al., 2024), these countervailing effects did not offset cattle impacts here.

Longevity declined with higher pressure and with more recent use: median survival fell from 460 to 256 days under high versus moderate pressure, and from 460 to 199 days in recently grazed versus ‘old’ (two-year hiatus) paddocks. These main effects fit herbivory theory—frequent defoliation and trampling reduce photosynthetic area, exhaust reserves, and delay ‘escape in size’ (Belsky et al., 1993; Rosenthal & Kotanen, 1994)—while compaction and litter removal depress moisture and organic matter, further limiting recovery (Pulido et al., 2010; Laskurain et al., 2013; Moradi et al., 2021). At paddock scale, intense or continuous use reduces safe sites and regeneration nuclei, narrowing the window for persistence despite any competition release from grazed swards (Tyler et al., 2008; Muñoz et al., 2009; López-Sánchez et al., 2016; Perea et al., 2017).

The pressure × history interaction indicates that a two-year cattle absence boosts survival only where grazing pressure was moderate. Under such conditions, release from cattle plausibly enables recovery of photosynthetic tissue, rebuilding of carbohydrate reserves, and root consolidation before renewed disturbance. Under high pressure, however, two years appear insufficient to reverse cumulative defoliation and compaction, an ecological legacy of heavily used pastures (Aronson et al., 2009; Pulido et al., 2010). Thus, regeneration is both intensity-dependent and temporally contingent: cohorts require moderate use plus extended rest to pass the most vulnerable stages.

Seedling height strongly predicted persistence (a 15 cm seedling lived 58% longer than a 5 cm seedling), consistent with greater photosynthetic tissue, biomass, deeper roots, and reserves buffering stress and aiding recovery (Belsky et al., 1993; Rosenthal & Kotanen, 1994; Kobe et al., 1995; Tyler et al., 2006; Leiva & Díaz-Maqueda 2016). Grazing suppresses height via browsing and trampling and indirectly through compaction and litter removal that reduce moisture and organic matter (Marcora et al., 2013; Laskurain et al., 2013; Moradi et al., 2021; Amsten et al., 2020). Because cattle target small, palatable shoots with limited structural or chemical defenses (Carmona et al., 2013; Göldel et al., 2016; Roula et al., 2019), intermittent escape in size can improve survival under continued use (Uytvanck et al., 2010). This also supports the management principle of allowing preferred woody species to grow beyond the reach of animals before renewed grazing (Öllerer et al., 2019).

Successful resprouting after top-kill was the strongest predictor of persistence. After top-kill, cork oak seedlings can rebuild canopy area from root reserves and re-mobilization of non-structural carbohydrate (NSC) reserves, a classic plant tolerance pathway under recurrent herbivory (Belsky et al., 1993; Rosenthal & Kotanen, 1994; Fornoni, 2011; Clarke et al., 2013; Pausas et al., 2018; Smith et al., 2018; Resco de Dios et al., 2020). In *Quercus*, resprouting vigor scales with root NSC pools and below-ground bud banks, supported by high root:shoot allocation and plastic hydraulic adjustments sustaining regrowth under drought (Clarke et al., 2013; Smith et al., 2018; Resco de Dios et al., 2020; Pausas et al., 2018). In cork oak, early taproot development and cotyledonary reserves fuel bud activation and shoot replacement, promoting persistence through summer droughts and after damage (Arosa et al., 2015; Zhang et al., 2019; Petersson et al., 2020). Seedlings that successfully resprouted persisted far longer than seedlings not observed to resprout, consistent with the role of protected buds and below-ground reserves in recovery after top-kill.

Shrub proximity conferred a modest advantage (∼6% longer predicted lifespan beneath shrubs vs. 5 m away), consistent with nurse-plant effects that buffer heat and water stress, improve soil infiltration, and reduce trampling/browsing (Gómez-Aparicio et al., 2004; Smit et al., 2008; Plieninger et al., 2004; Pulido & Díaz, 2005; Díaz et al., 2021). In our system, *Ulex* spp. likely supplied shade, physical protection, and soil N inputs, favouring early cork oak seedling establishment (Gómez-Aparicio et al., 2004; Duponnois et al., 2011; Dias et al., 2016). Other shrubs can act as competitors, however, so net effects depend on species identity and management. Because shrub clearing is routinely used to maintain open structure and reduce fire risk, the consequences for recruitment will differ if nurse shrubs are removed versus primarily competitive ones (Aronson et al., 2009; López-Sánchez et al., 2016). Prior work shows shrub cover can partially offset grazing impacts on oak seedlings (Pulido et al., 2010; Plieninger et al., 2011; Perea et al., 2016; Mochi et al., 2022); in our study this facilitation was detectable but small relative to the influences of grazing regime and resprouting.

Local dung abundance, a proxy for recent cattle activity, was positively associated with survival. This likely reflects fine-scale heterogeneity in cattle use: animals concentrate in microsites with better shade, moisture, or forage that also favor seedlings (Ganskopp & Bohnert, 2009). Dung can locally enrich organic matter and nutrients, enhance water-holding capacity, and stimulate bioturbation (Huerta et al., 2018; Wang & Hou, 2023), sometimes offering short-term protection from trampling/browsing (Ocumpaugh et al., 1996) and reducing desiccation risk (Leiva & Sobrino-Mengual, 2023). Dung/urine patches may also reduce local defoliation through fouling, because cattle often avoid feeding immediately adjacent to fresh excreta; this may indirectly lower herbivory risk for nearby seedlings (Whistance et al., 2011; Seó et al., 2015). Such signatures can create microsites of higher survival within broadly negative grazing contexts.

Wetter recent conditions (higher 1-month SPEI) were also associated with longer survival, consistent with water-balance controls on Mediterranean recruitment: brief improvements in hydraulic status can lift persistence during vulnerable stages (Vicente-Serrano et al., 2010; Beguería & Vicente-Serrano, 2023). Although cork oak exhibits drought-coping traits—early taproot development and below-ground reserves (Costa et al., 2010; Mendes et al., 2016; Arosa et al., 2015)—recent climate still mattered. Thus, while grazing history, pressure, and resprouting dominated, short wet spells provided a consistent, positive nudge to longevity.

### 4.2 Grazing and probability of seedling resprout

Cattle grazing reduced the likelihood that seedlings would resprout, with the greatest negative effect observed under recent use. Mechanistically, successful resprouting after top-kill requires an intact bud bank at the root collar/lignotuber and sufficient below-ground carbohydrates to fuel rapid shoot replacement; frequent, recent defoliation and trampling damage basal meristems and draw down reserves, suppressing bud activation and sprout persistence—classic tolerance dynamics under recurrent disturbance (Belsky et al., 1993; Rosenthal & Kotanen, 1994; Fornoni, 2011; Clarke et al., 2013). Two proximate levers align with our pattern: (i) reserve status and stubble integrity—seedlings retaining cotyledons and adequate residual shoot tissue resprout more vigorously (Arosa et al., 2015; Zhang et al., 2019; Petersson et al., 2020); and (ii) hydraulic safety—brief rest periods can restore water status and permit reserve rebuilding, whereas late-season, repeated use pushes plants toward hydraulic failure, limiting sprout initiation (Clarke et al., 2013). Accordingly, a two-year release of grazing in “old” paddocks likely allowed reserve recovery and bud maintenance, whereas “recent” grazing curtailed both processes, lowering resprouting odds.

### 4.3 Management implications

Our results point to simple, testable rules for keeping recruitment compatible with production in cork oak wood-pastures. First, stocking must be moderated and, critically, interspersed with multi-year rests. A two-year cattle absence improved survival only where grazing pressure was moderate; under high pressure it was insufficient—evidence of legacies that require longer recovery windows. Managers should therefore scale rest duration to recent pressure (≥2 years under moderate use; longer where high pressure has been chronic) and avoid repeated “recent” use on the same paddocks across consecutive years. Aligning rest periods with wetter spells further boosts survival, suggesting rotations that anticipate near-term water balance (e.g., SPEI-informed moves) can materially raise recruitment odds. However, this should not be interpreted as advocating grazing immediately after rainfall, when wet soils may increase uprooting and trampling damage.

Second, because resprouting is the dominant axis of persistence yet is itself depressed by recent grazing, management should explicitly protect the bud bank. Practical levers include: (i) preventing late-season, repeat use that compounds top-kill and basal damage; (ii) prioritizing paddocks or lighter stocking in paddocks holding many small seedlings; and (iii) using seedling height as a simple field indicator of recruitment—once cohorts gain height, subsequent mortality risk drops. Shrub cover offers modest facilitation; selective retention of nurse shrubs (e.g., *Ulex* patches) in recruitment zones can complement grazing rests, while indiscriminate clearing may erase these refuges. Dung patches flagged microsites with higher persistence; they can reveal fine-scale heterogeneity that can guide where to place micro-exclosures or adjust animal distribution. Together, these actions link our results—grazing pressure, grazing history, resprouting, and short-term climate—to practical grazing rotations that balance livestock use and tree regeneration in Mediterranean oak agroforestry.

## 5. Conclusion

Our study shows that cork oak recruitment in Mediterranean wood-pastures depends strongly on how cattle use is distributed through time and intensity. Recent grazing and high grazing pressure reduced both survival and resprouting, whereas moderate use combined with sufficiently long rest periods improved the prospects for persistence and recovery. These findings support adaptive grazing rotations that protect seedling cohorts through vulnerable stages and better align livestock production with oak regeneration.

## CRediT authorship contribution statement

**Abdullah Ibne Wadud**: Data curation, Formal analysis, Investigation, Methodology, Visualization, Writing – original draft. **João Craveiro**: Investigation, Writing – review & editing. **Simone Erroi**: Investigation, Writing – review & editing. **Sandra Alcobia**: Data curation, Writing – review & editing. **Miguel N Bugalho**: Supervision, Writing – review & editing. **Manuela Branco**: Conceptualization, Supervision, Writing – review & editing. **Pedro Gonçalves Vaz**: Conceptualization, Supervision, Methodology, Resources, Formal analysis, Visualization, Writing – review & editing.

## Acknowledgements

AIW (PT/BD/143139/2018) was funded by the Portuguese Science and Technology Foundation (FCT) through SUSFOR—Sustainable Forests and Products Doctoral Program (PD.00157.2012) and institutional supported by CEABN-InBIO UIDB/50027/2025; InBIO DOI 10.54499/LA/P/0048/2020). JC and PGV were funded by FCT through CE3C (UID/00329/2025) and CHANGE (DOI 10.54499/LA/P/0121/2020). PGV received funding from the AdaptForGrazing project (PRR-C05-i03-I-000035-LA4.3/4.4/4.6/4.7), supported by the EU (RRF) via IFAP (PRR).

## Notes

### Competing Interest Statement

The authors have declared no competing interest.

### Summary of Updates

This version includes suggestions from reviewers and should be close to the final version of the article to be published

