## Appendix A for "Recent and high grazing pressure limit cork oak seedling resprouting and survival"

**Table A.1.** Summary of the fixed effects from the mixed-effects Bayesian models evaluating the impact of cattle grazing on cork oak seedling survival. The top section presents results from a model comparing grazed and ungrazed paddocks, while the bottom section refers to grazed paddocks only and evaluates the effects of grazing pressure (moderate vs. high), grazing history (old vs. recent), and additional covariates. CI = 95% credible interval; β<>0 = posterior probability that the effect is greater (or smaller) than zero, depending on the sign of the estimate. The potential scale reduction factor (Rhat) was 1.00 for all parameters.

| *Model* / Parameter | *β* | Error | 2.5% CI | 97.5% CI | *β*<>0 |
| --- | --- | --- | --- | --- | --- |
| *Grazed vs. ungrazed* |  |  |  |  |  |
| Intercept | 6.20 | 0.15 | 5.89 | 6.48 | 1.00 |
| Grazed (yes) | -0.48 | 0.22 | -0.90 | -0.05 | 0.98 |
| *Grazing pressure & history* |  |  |  |  |  |
| Intercept | 5.89 | 0.23 | 5.45 | 6.38 | 1.00 |
| Grazing pressure (high) | -0.59 | 0.32 | -1.25 | 0.03 | 0.97 |
| Grazing history (recent) | -0.85 | 0.33 | -1.52 | -0.25 | 1.00 |
| Seedling height | 0.15 | 0.01 | 0.13 | 0.17 | 1.00 |
| Resprout (yes) | 1.63 | 0.06 | 1.52 | 1.74 | 1.00 |
| Distance to shrub | -0.02 | 0.01 | -0.05 | 0.00 | 0.95 |
| No. of dung | 0.44 | 0.04 | 0.37 | 0.51 | 1.00 |
| SPEI | 0.58 | 0.01 | 0.56 | 0.60 | 1.00 |
| Grazing pressure (high) × Grazing history (recent) | 1.22 | 0.45 | 0.35 | 2.15 | 1.00 |

**Table A.2.** Summary of the fixed effects from the mixed-effects Bayesian logistic regression models evaluating the effects of cattle grazing on the probability of cork oak seedling resprouting. The top section compares grazed and ungrazed paddocks; the bottom section refers to grazed paddocks only, evaluating the effects of grazing pressure and grazing history. Model output structure and notation are as in Table A.1.

| *Model* / Parameter | *β* | Error | 2.5% CI | 97.5% CI | *β*<>0 |
| --- | --- | --- | --- | --- | --- |
| *Grazed vs. ungrazed* |  |  |  |  |  |
| Intercept | -1.44 | 0.25 | -1.94 | -0.97 | 1.00 |
| Grazed (yes) | -0.78 | 0.39 | -1.60 | -0.05 | 0.98 |
| *Grazing pressure & history* |  |  |  |  |  |
| Intercept | -1.63 | 0.61 | -2.83 | -0.40 | 0.99 |
| Grazing pressure (high) | -0.79 | 0.73 | -2.32 | 0.58 | 0.87 |
| Grazing history (recent) | -1.56 | 0.71 | -3.03 | -0.20 | 0.99 |
